# A Dual-Action Liposome-Peptide Formulation Synergistically Counteracts A Gain-of-Function p53 Mutant

**DOI:** 10.1101/2025.04.09.647913

**Authors:** Sneha Ghosh Chaudhary, Swati Bhowmick, Samriddhi Bhattacharya, Siddhartha Roy, Nahid Ali

**Affiliations:** CSIR-IICB, 4, Raja S. C. Mullick Road, Kolkata-700032; Bose Institute, EN 80, Sector V, Bidhan Nagar, Kolkata - 700091; Experimental Immunology Branch, National Cancer Institute, National Institute of Health, USA

**Keywords:** mutant p53, wild-type p53, cationic liposome, chemoresistance, gain-of-function

## Abstract

**Objective:** Inactivation of p53 tumor suppressor functions, often through missense mutations, is essential for carcinogenesis. A sub-class of such p53 missense mutations gains new functions, including drug resistance and enhanced proliferation, in addition to its loss of function. Among the most frequent gain-of-function p53 mutants, R273H occurs in tumors of many tissue origins and imparts aggressive character and resistance to drugs to the tumor. Tumors bearing p53R273H are generally resistant to all available therapies, and need for novel interventions are urgently needed. Interaction of p53R273H with Positive Coactivator 4 (PC4), an abundant chromatin-associated protein, is essential for acquiring the gain-of-function properties. Previously, we developed a chemically modified peptide, NLS-p53(380-386), targeting PC4 that abrogated the interaction of p53R273H with PC4 and reversed many of its gain-of-function properties. We earlier demonstrated that cationic phosphatidylcholine-stearylamine (PC-SA) liposomes possess inherent anti-tumor properties. To improve efficacy, pharmacokinetics, and delivery, we entrapped the PC4-targeted peptide into PC-SA liposome.

**Methods:** We synthesized the NLS-p53(380-386) peptide and entrapped in PC-SA liposome. We used MTT assay, confocal microscopy, flow cytometry, qRT-PCR, and western blotting to investigate the biological effects of the p53-entrapped PC-SA.

**Results:** Pre-treatment with the PC-SA liposome entrapped peptide enhanced the chemosensitivity of widely used anticancer drug doxorubicin in cell lines bearing p53R273H mutation. The doxorubicin-induced cell-killing effect was much more enhanced when pre-treated with the liposome-entrapped peptide than when pre-treated with either the free peptide or the liposome alone.

**Conclusion:** The liposome-encapsulated peptide is a promising formulation for developing therapies targeting tumors bearing the p53R273H.

## Introduction

The tumor suppressor protein p53, encoded by the gene TP53, is critical for maintaining cellular homeostasis and protecting cells from tumorigenesis (1), (2), (3). It is well established that more than half of all types of human cancers contain altered TP53 alleles that lose their tumor suppression properties, frequently through missense mutations (3). Most of the mutations in the TP53 gene are missense mutations that cause single amino acid substitutions in the wild-type protein. Although p53 mutations occur in all coding exons of the protein, the majority of p53 mutations occur in the DNA-binding domain (DBD), resulting in a deficiency of binding to cis-regulatory elements of its target genes and therefore impaired transcriptional regulation of those genes. Approximately, one-third of all p53 mutations occur at the six hotspot residues in the DBD region, Arg-175(R175), Gly-245(G245), Arg-248(R248), Arg-249(R249), Arg-273(R273), Arg-282(R282) (2), (4). While the majority of the mutant p53 proteins lose their tumor-suppressive functions and show dominant-negative effects over the wild-type p53, some mutants such as R248Q, R273H, R175H, and R249S, acquire new growth-promoting functions, called ‘gain-of-function (GOF)’, that act as drivers of aggressive growth properties, metastasis and chemoresistance (4). Since p53 GOF mutants are often associated with late-stage malignancy and drug resistance, novel therapies targeting such hotspot p53 mutants may address an area of urgent need (3), (4), (5), (6), (7).

p53 interacts with many cellular proteins. These interactions underpin most of its functions. Positive cofactor 4 (PC4) also interacts with p53, enhancing p53 binding with cis-regulatory elements, resulting in augmented transcription of p53-mediated downstream genes and apoptosis (8), (9). Residues (380-386) of the p53 protein were found to be crucial for binding with PC4 and p53-PC4 interaction was further enhanced by the acetylation of lysine residues 381 and 382 (10). In cells harboring p53R273H mutation-such as SW480 and PANC-1-the p53 mutant piggybacks on pro-oncogenic transcription factors to relocate to the promoters of pro-oncogenic genes despite the loss of its DNA-binding activity. Previously, we have shown that recruitment to some of these promoters occurs by the formation of a stable protein complex with PC4. It was also shown that PepAc disrupted the binding of the wild-type p53 with PC4 (10). PepAc also disrupted the binding of a GOF-mutp53, p53R273H with PC4, and the application of RN-PepAc to p53R273H harboring cells resulted in the reversal of many Gain-of-function properties of the p53R273H protein (11). This study by Mondal et. al successfully established that RN-PepAc disrupts the interaction of PC4 and p53R273H, inhibiting the recruitment of the mutant p53-containing complex on the promoters like MDRI and UBEC2. The promise of these observations for development of new therapies is tempered by the fact that peptides suffer from limitations in *in vivo* applications, due to their shorter half-life in body fluids, poor cell permeability, and higher rates of enzymatic degradation, thus requiring an appropriate delivery system for *in vivo* applications (12), (13), (14).

Tissue-targeted delivery of chemotherapeutic agents that spare healthy tissues remains a challenge in drug development. Several drug delivery vehicles like liposomes, nanotubes, and nanostructured liquid crystalline particles such as cubosomes or hexasomes have made significant advances in targeted drug delivery in cancer and other diseases (15), (16). We previously reported the preparation and use of phosphatidylcholine-stearylamine (PC-SA) liposomes, which can target tumor cells specifically as SA has a strong affinity and direct interaction with the exposed phosphatidylserine (PS) on the tumor cells (17). The same report also showed that PC-SA liposomes by themselves possess anti-tumor properties and entrapment of common anticancer drugs enhanced their therapeutic efficacy many folds, both *in vitro* and *in vivo,* without any noticeable toxicity.

In this study, we entrapped the peptide N-PepAc in PC-SA liposomes-from hereon will be called PC-SA:N-PepAc and targeted cancer cells carrying the R273H mutation in their TP53 gene. We demonstrated that PC-SA:N-PepAc enhanced the chemosensitivity of the cell lines bearing the p53R273H to doxorubicin (DOX) at much lower peptide concentrations than the equivalent RN-PepAc. This low dose of the peptide was enough to bring about the abrogation of activation of several p53R273H target genes like the multidrug-resistant gene MDR1, proliferating cell nuclear antigen (PCNA), apoptosis-inducing genes, signaling caspases and reduction of reactive oxygen species (ROS). This indicates that PC-SA:N-PepAc could be an effective formulation for selective targeting of p53R273H protein and potentially other GOF p53 mutants. It may offer a way to efficacious anti-tumor therapy acting through two different mechanisms (11), (17).

## Methods

### Reagents, Kits and Antibodies

Fmoc-amino acids, Rink amide MBHA (4-methylbenzhydrylamine) resin; TBTU, O-(Benzotriazol-1-yl)-N,N,N’,N’-tetramethyluronium tetrafluoroborate; HOBT, 1-Hydroxybenzotriazole hydrate (Novabiochem, Mumbai, India); DMF, Dimethylformamide, Piperidine, DIPC, N,N′-Diisopropylcarbodiimide; DIPEA, N,N-Diisopropylethylamine; HPLC-water, acetonitrile, TFA (Spectrochem, Mumbai, India), Reverse-phase C-18 HPLC column (Thermo Corporation Limited, Mumbai, India); Fluorescein isothiocyanate (5,6-carboxy fluorescein) (FITC), propidium iodide (PI) (Invitrogen, Bangalore, India); Dulbecco’s Modified Eagle’s Medium (DMEM), Fetal bovine serum (FBS), Penicillin-Streptomycin (10,000 U/mL), Trypsin-EDTA (0.5%) (Gibco-BRL, Gaithersburg, USA); DOX, 3-(4, 5-dimethylthiazolyl-2)-2, 5-diphenyltetrazolium bromide (MTT), Primers (Sigma-Aldrich, Bangalore, India), RNeasy Plus (Qiagen Inc., CA, USA), DreamTaq^TM^ Green PCR Master Mix (Thermo Fisher Scientific, Rockford, USA); iScript cDNA Synthesis Kit (Bio-Rad, Haryana, India); 2X DyNAmo Color Flash SYBR Green Master Mix (Thermo Fisher Scientific); BCA protein assay kit (Thermo Fisher Scientific); FITC Annexin V/Dead Cell Apoptosis Kit (eBioscience, San Diego, CA); Cleaved Caspase Antibody Sampler Kit (Cell Signaling Technology, Danvar, USA), Primary antibodies NOXA, PUMA and BAX (Cell Signaling Technology); egg PC (Sigma Aldrich), SA (Fluka, Buchs, Switzerland)

### Peptide Synthesis and Purification

The sequences of the synthesized peptides are listed in Table 1. The peptides were synthesized on a 0.1-mmol scale by solid phase peptide synthesis strategy using 9-fluorenylmethoxy carbonyl chemistry and Rink amide MBHA resin (substitution 0.36 mmol/g) as described before (18), (19).

**Table 1:** Sequence of the peptides synthesized.

| Peptides synthesized | Sequence |
| --- | --- |
| NLS-p53(380-386) or N-PepAc | PKKKRKVHKAcKAcLMFKAc |
| R6-NLS-p53(380-386) or R-N-PepAc | RRRRRR PKKKRKVHKAcKAcLMFKAc |

KAc stands for acetylated residues, the changes in control peptides are shaded and R6 indicate six consecutive D-arginine.

Briefly, the synthesis procedure includes swelling and deprotection of resin, coupling of the amino acids to the resin, and finally cleavage of the peptide chain from the resin. The entire procedure of the solid support peptide synthesis was performed in an inert nitrogen atmosphere. Required amount of weighed resin in a clean and dry peptide synthesizer vessel was swelled in DMF for an hour and deprotected with 8% piperidine (v/v) in DMF to yield a free –NH_2_ group for subsequent addition of amino acids. To test the proper decoupling, a small quantity of the resin from the peptide synthesizer column was allowed to react for 1 min with an aliquot of trinitrobenzene sulfonic acid (TNBS)/picrylsulfonic acid solution, 5% (w/v). A bright reddish orange color of the bead indicates the presence of free amino group available for coupling. The deprotected resin was allowed to couple with the amino acid dissolved in DMF with TBTU as a peptide coupling agent and DIPEA as a base for 40-45 min under inert nitrogen atmosphere. After thorough washing to remove uncoupled amino acids, successive deprotection and coupling continued to synthesize the peptide.

The synthesized peptide was cleaved from the resin by a cocktail containing TFA (87.5%), EDT (2.5%), methylphenyl sulfide/thioanisole (5%), dimethyl sulfide (2%), HPLC grade water (3%) and ammonium iodide (1.5% w/v) followed by precipitation in cold diethyl ether. The precipitate was collected by centrifugation, dried, and dissolved in 0.1% TFA water.

A small portion of the synthesized peptide was labelled at the N-terminus with FITC in solid phase. The resin was first washed with 20% piperidine and the resin bound N-terminal deprotected peptide was reacted with 10 times molar excess of FITC, HOBt and DIPC at 25 °C for 3 h. The labelled peptide was cleaved from the resin as described above.

The crude peptide was purified by reversed-phase high performance liquid chromatography (HPLC) on a C18 column with linear gradients of 0-80% water/acetonitrile containing 0.1% trifluoroacetic acid. Peptide masses and purity (>95%) were checked by MALDI-TOF Mass Spectrometry.

### Entrapment of peptide in liposome and characterization

Liposome containing peptide was prepared as described earlier with slight modifications (17),

(20). Lipids PC-SA (7:2 molar ratio) were dissolved completely in chloroform and a thin film was produced by evaporating the organic solvent by rotation in a round-bottom flask and dried overnight in a vacuum desiccator. Next day the film was rehydrated with 500 µL of 20 mM PBS (pH 7.4) containing peptide (300 µg peptide/20 mg lipid) and sonicated in an ultrasonicator (Misonix, Microson; output energy, 4 W) for 60 s. Unentrapped free peptide was removed by centrifugation at 100,000 x g for 1 h at 4°C. The amount of peptide entrapment was estimated by BCA protein assay kit.

Free and peptide entrapped PC-SA liposome size distribution (measurement of hydrodynamic diameter) and zeta potential were analyzed by laser DLS (Malvern Instruments, Zeta sizer, Nano-ZS, model ZEN 3600).

### Cell Culture

Human colorectal adenocarcinoma cell line SW480 and human pancreatic duct epithelioid carcinoma cell line PANC-1 were kind gifts from Centre for Cellular & Molecular Biology, Hyderabad, India. HEK293 cells were a gift from a lab at CSIR-IICB. Cells were sub-cultured in complete medium containing DMEM, 10% FBS and penicillin/streptomycin in the presence of 5% CO_2_ at 37 ^0^C.

Morphology of PC-SA entrapped NLS-p53(380–386) was studied by AFM (Bruker, USA). Liposomes dispersed in Milli-Q water (1:100) were spotted on freshly cleaved mica sheets and dried to remove water. The dried sheets were subjected to AFM analysis.

### Analysis of PS exposure

PS exposure of the cell lines was analyzed by FITC Annexin V staining. Cells were harvested and washed with cold PBS. Cell pellets were suspended in 1 X Annexin-V binding buffer at a concentration 10^6^ cells/mL. 100 µL of cell suspension was stained with 5 µL annexin V-FITC at room temperature for 15 min in the dark. PS analysis was carried out in flow cytometer (BDLSRFortessa^TM^ Cell Analyser, BD).

### Confocal Microscopy

To check the entry of peptide into cell and subsequent entry into nucleus for interaction with DNA, PANC-1 cells (10^5^ cells/10 mm confocal plate) were incubated with liposome containing FITC-labelled peptide for 2 h, followed by addition of fresh medium and kept for 24 h. At 2 h and 24 h time points, cells were fixed with 4% paraformaldehyde for 20 min, washed with PBS and stained with Hoechst (nucleus stain). The culture plates were observed under Leica Stellaris5 confocal microscope (FITC excitation and emission wavelength was 513 nm and 565 nm respectively, Hoechst excitation and emission wavelength were 429 nm and 513 nm respectively (21).

### MTT assay

Viability of cells with graded doses of free DOX, free PC-SA, free peptide, and liposomal peptide were determined by MTT assay. Freshly harvested cells were seeded in a 96 well cell culture plate at a density of 5 × 10^4^ cells/well followed by different treatment conditions. For free DOX, cells were incubated with 0 to 1.0 µg/mL of DOX and kept for 72 h. For free PC-SA, cells were incubated with graded doses of liposome (0 to 140 µg/mL) for 2 h, fresh media was added to remove excess free liposome and kept for 72 h. At 72 h, the cells were centrifuged, washed with 20 mM PBS and incubated with 2 mg/mL MTT solution for 2 h at 37 C. DMSO was added to dissolve the formazan crystals and absorbance was recorded on an ELISA plate reader (Thermo) at 550 nm. EC_50_ concentration was determined by fitting to an EC_50_ function for analysis of the dose response values.

For peptides, the cells were incubated with 0 to 100 µg/mL N-PepAc or R-N-PepAc, followed by addition of 0.5 µg/mL dose of free DOX at 24 h and kept for 72 h. For the treatment with liposomal peptide, cells were incubated with 40 to 100 µg/mL PC-SA peptide for 2 h followed by fresh media change, kept overnight and addition of 0.5 µg/mL dose of free DOX at 24 h and left for 72 h. MTT addition and readings were taken by the same method described above at 72 h.

To assess the caspase-dependent cell death, cells were preincubated with pancaspase inhibitor Z-VAD-FMK (Sigma) followed by incubation with free DOX (0.5 µg/mL) or graded doses of PC-SA-N-PepAc. The treatment protocol for the two groups were same as described above.

### Apoptosis assay

SW480 cells (1 x 10^6^) were seeded in 6-well plates and incubated with only media (control), 0.5 µg/mL dose of free DOX, 60 µg/mL PC-SA with and without N-PepAc and N-PepAc or R-N-PepAc (corresponding dose entrapped in liposome). For liposome treatment, fresh media was added after 2 h as described above. Following DOX addition at 24 h, cells were harvested for 48 h time point and stained with Annexin V-FITC Apoptosis Detection kit according to manufacturer’s instruction. Apoptosis analysis was carried out on BDLSRFortessa^TM^ Cell Analyzer, BD.

### qRT-PCR

SW480 cells were plated in 6 well plates in three groups. In one group, cells were incubated with PC-SA or PC-SA-N-PepAc for 2 h, or 1µg/ml of free peptide for 24 h. Excess liposomes were washed off after 2 h and fresh media added. Next day, 0.5 µg/ml of free DOX was added and kept for another 24 h. Cells without treatment and treated with only DOX 24 h or only peptide and only liposomes entrapped peptide (without DOX) were the other groups kept as controls. RNA was isolated from control or treatment groups with RNeasy Plus and concentration was measured using Nanodrop 2000 (Thermo Scientific, USA). Approximately, 2 μg RNA was reverse transcribed using iScript cDNA synthesis kit in 20 μL reaction volume: priming 25 C for 5 min, reverse transcription at 46 C for 20 min and heat inactivation at 95 C for 1 min.

To determine quantitative mRNA expression of the respective genes, real time RT-PCR was carried out in Roche Light Cycler 96 using 10 μL reaction mixture containing 2 μL cDNA, 0.5 μL primer pairs (25nM), 5 μL 2X DyNAmo Color Flash SYBR Green Master Mix and nuclease-free water. The primer sequence, annealing temperature and product size of all the genes are listed in Table 2. For RT-PCR, 2 μL cDNA, 0.5 μL 25 nM primer pairs of respective gene, 5 μL 2X DyNAmo Color Flash SYBR Green Master Mix and nuclease free water were added in 10 μL reaction mixture and PCR was performed in Roche Light Cycler 96. Ct values for control and test genes were analyzed using Light cycler 96 software and fold change was calculated as described earlier (21).

**Table 2:**
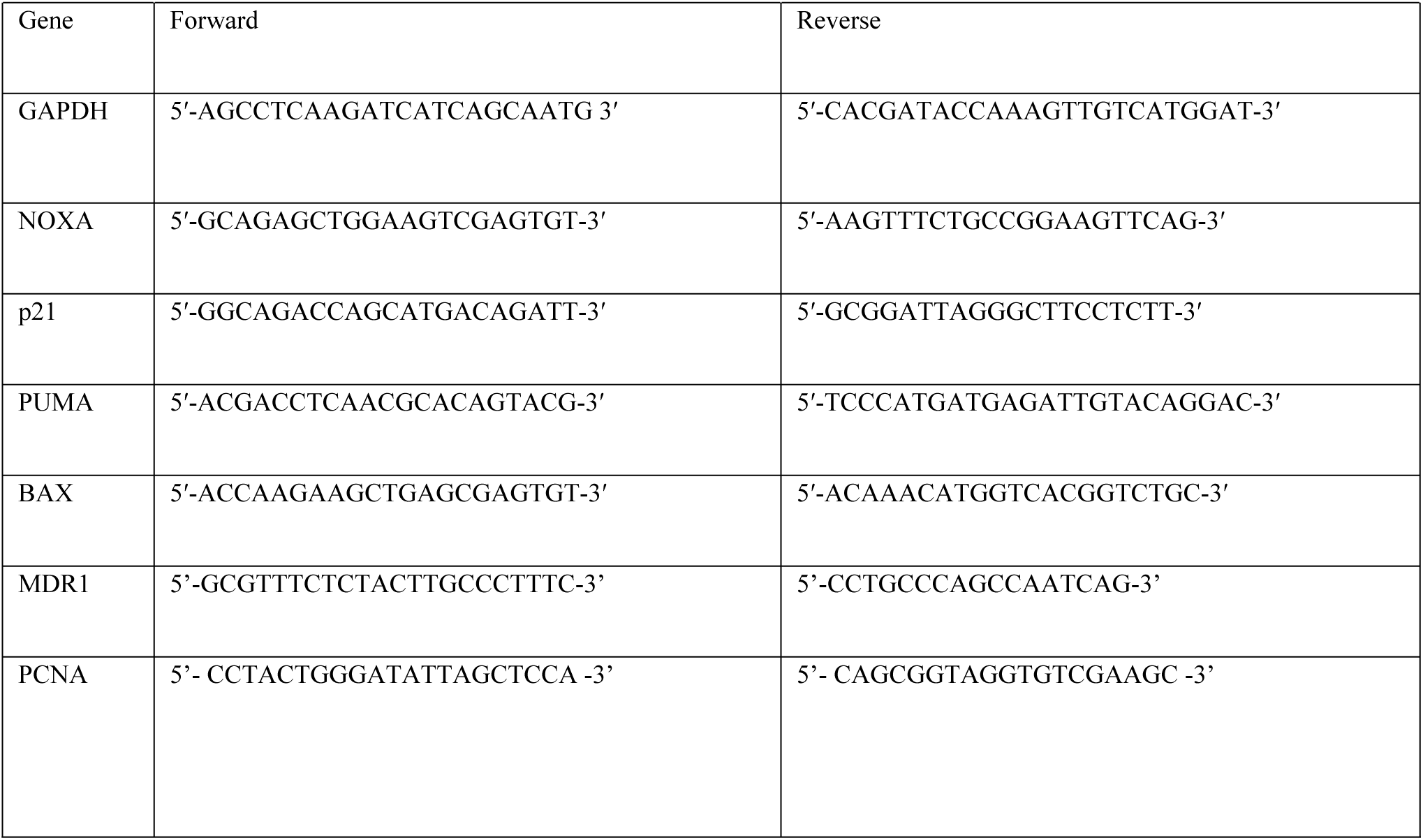
Primer sequences used for qRT-PCR.

### Western Blotting

Immunoblotting was carried out with the same three groups of cells with identical treatments as mentioned for RT-PCR studies. Following incubation period, cells were washed with PBS and lysed in cell lysis buffer (150 mM sodium chloride, 1.0% Triton X-100, 0.5% sodium deoxycholate, 0.1% sodium dodecyl sulfate (SDS), 10 mM Di-thiothreitol (DTT), 50 mM Tris, pH 8.0 containing protease and phosphatase inhibitor Cocktail (Roche, Germany). Bradford method was used to determine protein concentration in the cell lysates. After resolution on 10% SDS-PAGE the proteins were blotted on to nitrocellulose membranes (BioRad, US). Blots were developed using the following steps: blocking with 5% Bovine Serum Albumin (BSA) (Sigma Aldrich, US) in Tris-buffered saline (TBS) for 1 h at room temperature, incubating with primary antibodies overnight at 4 C, washing with wash buffer (TBS containing 0.05% tween), incubating with HRP-conjugated secondary antibodies (Cell Signaling Technology) and detection by Luminata Forte (Millipore Sigma, US) western HRP substrate (Fischer scientific, US).

### ROS generation

Intracellular ROS generation was estimated in cells after treatment with liposomal NLS-p53(380–386) using 2′,7′-dichlorodihydrofluorescein diacetate dye (Molecular Probes) as reported earlier (22). After 30 min of incubation, cells were washed in 20 mM PBS and analyzed by flow cytometry.

### Statistical Analysis

All statistical analysis was carried out using GraphPad Prism 9 software. Student’s t test was used to obtain statistical significance between two groups. *p* <0.05 was considered significant.

## Results

### The liposome entrapped peptide enters into nuclei of adenocarcinoma cells

It was earlier reported that the PepAc interacted with transcriptional coactivator PC4 with high affinity and when attached with NLS and cell penetration tags (R-N-PepAc) resulted in modulation of p53 regulated genes in cells (10), (11). This peptide also disrupted the interaction of R273H-p53 with PC4 resulting in the abrogation of many of its GOF properties in a cell (11). The unique ability of the stearylamine-bearing PC-SA liposome to bind to exposed PS of tumor cells has made it an attractive delivery agent for tumor-selective anti-cancer therapy (17). We have used PC-SA:N-PepAc for specifically targeting adenocarcinoma cells, to evaluate whether it is an effective formulation for further use *in vivo*. As compared to the previous study (11), the cell penetration tag on PepAc was omitted as the liposome should be able to deliver its cargo to the cytoplasm. The amount of peptide entrapment was 64% of starting amount. The PC-SA:N-PepAc was characterized by measuring its size, zeta potential, and observation of its morphology by AFM. Hydrodynamic diameter, as measured by laser dynamic light scattering (DLS) was found to be 273.1 ± 74.32 nm for free PC-SA liposomes and 366.7 ± 31.83 nm for PC-SA:N-PepAc with polydispersity index values less than 0.5 for both, thus indicating the systems had narrow size distribution with low aggregation of particles (Fig. 1A). Zeta potential of free PC-SA liposomes was +57.3 mV and that of the PC-SA:N-PepAc was +57.7 mV (Fig. 1B). To visualize the morphology of liposomes, an AFM analysis was performed. Topography flattened and amplitude flattened views of PC-SA:N-PepAc showed liposomes were clean spherical, well separated from each other. The heights of the liposomes were 4.5 ± 0.5 nm as displayed in the height profile (Fig. 1C).

**Fig. 1.**
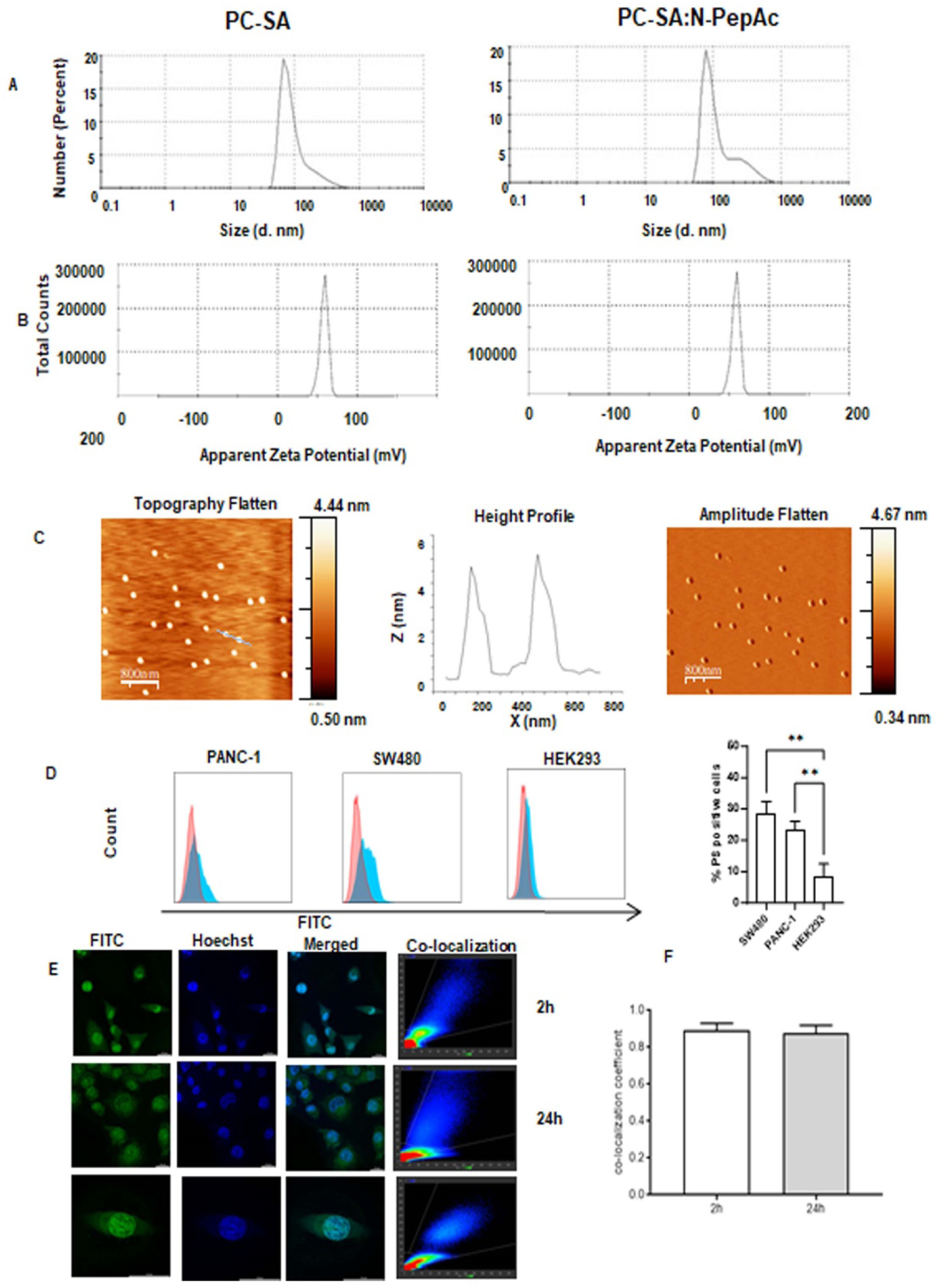
Characterization of the PC-SA:N-PepAc and intracellular localization of the peptide. **(A)** Size of the free PC-SA and PC-SA:N-PepAc. The image is a representative one, out of three independent measurements. **(B)** Zeta Potential distribution of the free PC-SA, and PC-SA:N-PepAc. The image is a representative one, out of three independent observations. **(C)** AFM images of PC-SA:N-PepAc are shown as topography-flattened, height profile, and amplitude flattened views. **(D)** Expression of PS on SW480, PANC-1 and HEK293 was assessed by FACS following Annexin V-FITC staining. The blue and red bars represent the unstained and stained populations, respectively. **(E)** Nuclear translocation of FITC labeled N-PepAc in PANC-1 cell line as observed by confocal microscopy. The bottom panel shows a single cell (enlarged view) with an arrow indicating the labeled peptide in the nucleus. All images were captured at 25 µm scale. **(F)** Co-localization coefficients of FITC signals with Hoechst signals were determined while capturing the images at 63X oil immersion objective with 3.0X optical zoom factor. (Significance is denoted by ** *p* < 0.01).

PC-SA liposomes target cancer cells by targeting exposed PS on the surface. We, therefore, assessed the expression of PS on two adenocarcinoma cell lines SW480 and PANC-1 which have p53R273H mutation. We found expression of a significant amount of PS on both the cell lines compared to normal cells like HEK (Fig. 1D). PC4 is a chromatin-associated protein, localized in the nucleus (23). We hypothesized that upon delivery to the cytosol, N-PepAc will migrate to the nucleus and bind to PC4 and disrupt the interaction with p53R273H. To verify this hypothesis, we labeled the N-PepAc with FITC and encapsulated it in the PC-SA liposomes and checked the entry of the peptide into the nucleus by confocal microscopy. As we incubated cells with PC-SA:N-PepAc for 2 h, we verified the cytosolic entry of N-PepAc and its migration to the nucleus at the end of 2 h and 24 h (Fig. 1E). At both the time points, merged images show significant co-localization of FITC signal with the signal from the Hoechst dye, a nuclear stain, which is clearly visible in the zoomed image of a single cell at the bottom panel. High co-localization coefficient values (>0.8) at 2 h and 24 h post-liposome treatment further suggested positive accumulation of the FITC-labeled N-PepAc in the nucleus of cells studied (Fig. 1F).

### The liposome-entrapped peptide enhanced drug sensitivity of cells carrying the p53R273H allele

Gain-of-Function p53 mutations are very frequent in various types of human cancer and contribute to enhanced growth and drug resistance. It is believed that GOF-mutantp53 acquires these additional functions through newly gained interactions with one or more pro-oncogenic transcription factors (24). In a previous study, it was shown that PC4 is also necessary for gain of these new functions by the GOF mutant p53R273H (25). Colorectal and pancreatic adenocarcinoma are two leading causes of worldwide cancer associated death. We used the colorectal cancer cell line SW480 and pancreatic cancer cell line PANC-1, both carrying the p53R273H hotspot mutation with associated GOF, including chemoresistance. It was hypothesized that N-PepAc delivered to cancer cells efficiently by PC-SA liposomes would abrogate GOF activity with higher efficacy, including chemoresistance, associated with this p53 mutation and make the cells more sensitive to DOX-induced death than the free peptide.

We first measured the EC_50_ doses of free DOX in SW480 and PANC-1 cell lines at 72 h, which were 0.54 µg/mL (1 µM) and 0.68 µg/mL (1.5 µM), respectively (Fig. 2A). As we had earlier observed anticancer activity of the PC-SA liposome alone, we determined the best suitable dose for peptide delivery by treating cells with graded doses of PC-SA for 2 h, followed by re-incubation in fresh media for 72 h (Fig. 2B). For SW480 and PANC-1, 60 µg/mL PC-SA showed 64% and 65% cell viability respectively, while higher doses were more cytotoxic. We next measured the effects of the free peptide on cell viability upon DOX treatment. This was performed by incubation of cells with graded doses of peptides, N-PepAc and R-N-PepAc (1-100 µg/mL) for 24 h, followed by treatment with 0.5 µg/mL dose of DOX (a dose close to EC_50_ fixed for both PANC-1 and SW480) for another 72 h. The effects are shown with the free peptides, N-PepAc (Fig. 2C) and R-N-PepAc, (Fig. 2D). At the lowest dose (1 µg/mL), the percentage of cell viability was comparable for both the peptides in two cell lines (46.66% and 45.66% in SW480 and PANC-1 respectively, for N-PepAc and 75% and 70% in SW480 and PANC-1 respectively for R-N-PepAc. At the highest dose (100 µg/mL), the N-PepAc showed 26.66% (SW480) and 20.3% (PANC-1) cell viability, whereas R-N-PepAc showed 10.63% (SW480) and 6.2% (PANC-1) viability, both upon DOX treatment for 72 h. This demonstrated the need for the peptide to cross the plasma membrane efficiently for it to be effective.

**Fig. 2.**
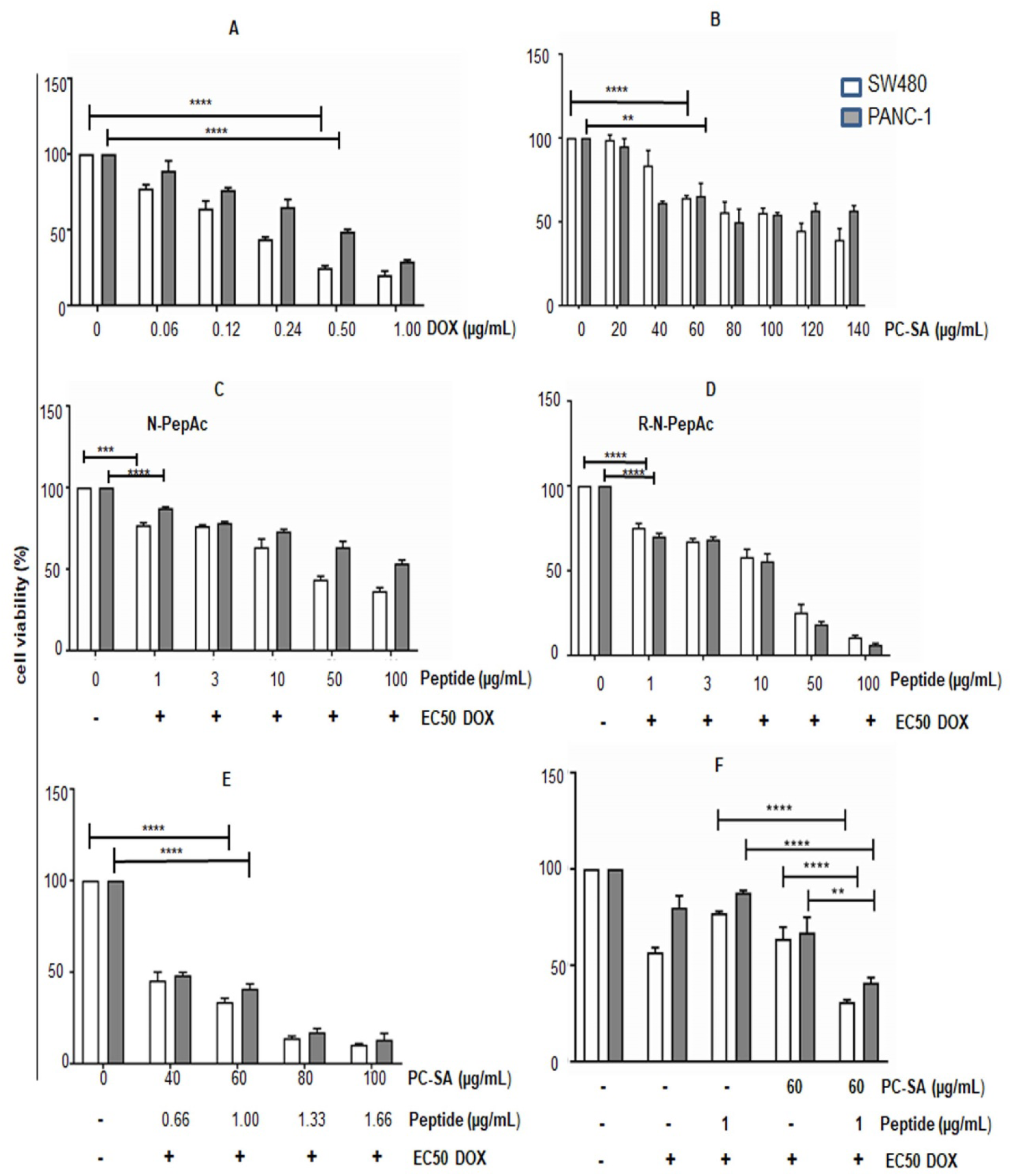
The PC-SA-N-PepAc increases the DOX sensitivity of cancer cells. **(A)** SW480 and PANC-1 cell lines were treated with graded doses of DOX for 72 h and analyzed by MTT. **(B)** Cell viability assay with graded doses of free PC-SA liposomes. **(C)** The effect of free N-PepAc on cancer cell lines to DOX treatment. The cells were treated with graded concentrations of peptide for 24 h followed by addition of EC_50_ dose of DOX and viable cell population at the end of 72 h was assayed. **(D)** The sensitization effect of free R-N-PepAc on both the cell lines to DOX treatment was observed in the same way as described for C. **(E)** The dose response of PC-SA:N-PepAc to EC_50_ dose of DOX. **(F)** Assessment of cancer cell viability on DOX treatment by 60 µg/mL of PC-SA liposome entrapping 1 µg N-PepAc and corresponding doses of free liposome and N-PepAc. All data are representative of three independent experiments. Significance is denoted by ** *p* < 0.01, *** *p* < 0.001, **** *p* < 0.0001.

Next, we evaluated the synergistic activity of the PC-SA:N-PepAc. For that, we incubated cells for 2 h with graded doses of PC-SA:N-PepAc followed by washing off the excess liposome and addition of DOX at 24 h and further incubation till 72 h. We found a significant dose-dependent decrease in cell viability by the PC-SA:N-PepAc with 60 µg/mL PC-SA carrying 1 µg/mL peptide reaching much-reduced viability in both SW480 (31.66%) and PANC-1 (38.01%) (Fig. 2E). Although increasing doses of the PC-SA:N-PepAc (80 and 100 µg/mL) induces even lower cell viability, a large number of cells were detached from cell plates and hence excluded from further investigation.

Notably, 1 µg/mL (0.5 µM) N-PepAc entrapped in 60 µg/mL PC-SA liposome induced almost the same DOX-induced cell death as we observed with the N-PepAc at 50 µg/mL (25 µM) (Fig. 2C), indicating that the efficiency of the peptide was enhanced very significantly when encapsulated in PC-SA liposomes. Although R-N-PepAc has a good ability to enter cells, it showed significant DOX-induced cell death only at higher doses. Thus, encapsulation of a peptide in PC-SA liposome could markedly reduce the effective dose of the peptide. As PC-SA liposome itself has anti-tumor properties, the higher cell death observed may be due to the combined activity of both the peptide and PC-SA liposome. To better understand the effectiveness of each component and the activity of their combination, if any, we evaluated cell viability following incubation with the selected dose of PC-SA:N-PepAc along with the equivalent doses of the N-PepAc, and liposome alone. In all cases, the dose of free DOX used was identical (Fig. 2F). SW480 cells incubated with PC-SA:N-PepAc demonstrated significantly higher sensitivity to DOX leading to lowered cell viability compared to either the N-PepAc (*P <*0.0001) or the free PC-SA (*P <* 0.0001). For PANC-1, combined activity was also significantly higher when compared to the N-PepAc (*P <*0.0001) and free PC-SA (*P <*0.05). Thus, a very low dose of N-PepAc when entrapped in PC-SA liposome could enhance sensitivity of mutant p53R273H-bearing cancer cells to DOX treatment and the effect was markedly better compared to monotherapies or without liposome entrapment.

### PC-SA entrapped peptide induced cell death by apoptosis following DOX treatment

Apoptosis induction is a goal of cancer treatment, and therapeutic strategies aim to induce or enhance the extent of apoptosis in cancer cells (26), (27), (28), (29), (30) (31). Here we checked whether the reduced cell viability, as determined by the MTT assay, was due to cell death and if so, by what mechanism it occurs following the DOX treatment. For that, we incubated SW480 cells with PC-SA:N-PepAc for 2 h, and then treated with EC_50_ dose of DOX at 24 h. PC-SA liposome without any peptide, N-PepAc and mock treatment along with DOX groups were kept for comparative evaluation. Cells were subjected to Annexin V-Propidium iodide (PI) staining at the end of 48 h post DOX-treatment. Cells negative for both PI and Annexin V (-PI, -Annexin, Q3) were considered live, negative for PI and positive for Annexin V (-PI, +Annexin, Q4) were considered early apoptotic, cells positive for both PI and Annexin V (+PI, +Annexin, Q2) were considered dead by late apoptosis, and cells positive for PI and negative for Annexin V (+PI, -Annexin, Q1) had undergone non-apoptotic death by necrosis. In all groups, large scale cell death was observed, suggesting that the major mechanism of reduction of cell viability was due to cell death. Flow cytometry-based dot plots in Fig. 3A demonstrate that cells treated with only DOX mostly died by necrosis. Incubation with free peptide (N-PepAc) resulted in both necrotic cell death and late apoptotic death in 48 h post DOX treatment. We observed lesser necrotic and more late apoptotic cell death by free PC-SA-treated group. For the group of cells treated with PC-SA:N-PepAc, a high percentage of late apoptotic cell death was observed in 48 h post DOX addition. Different stages of cell population percentage were represented as histograms in Fig. 3B. This switch in mechanism of cell death may indicate a more controlled action by DOX and lower off-target toxicity.

**Fig. 3.**
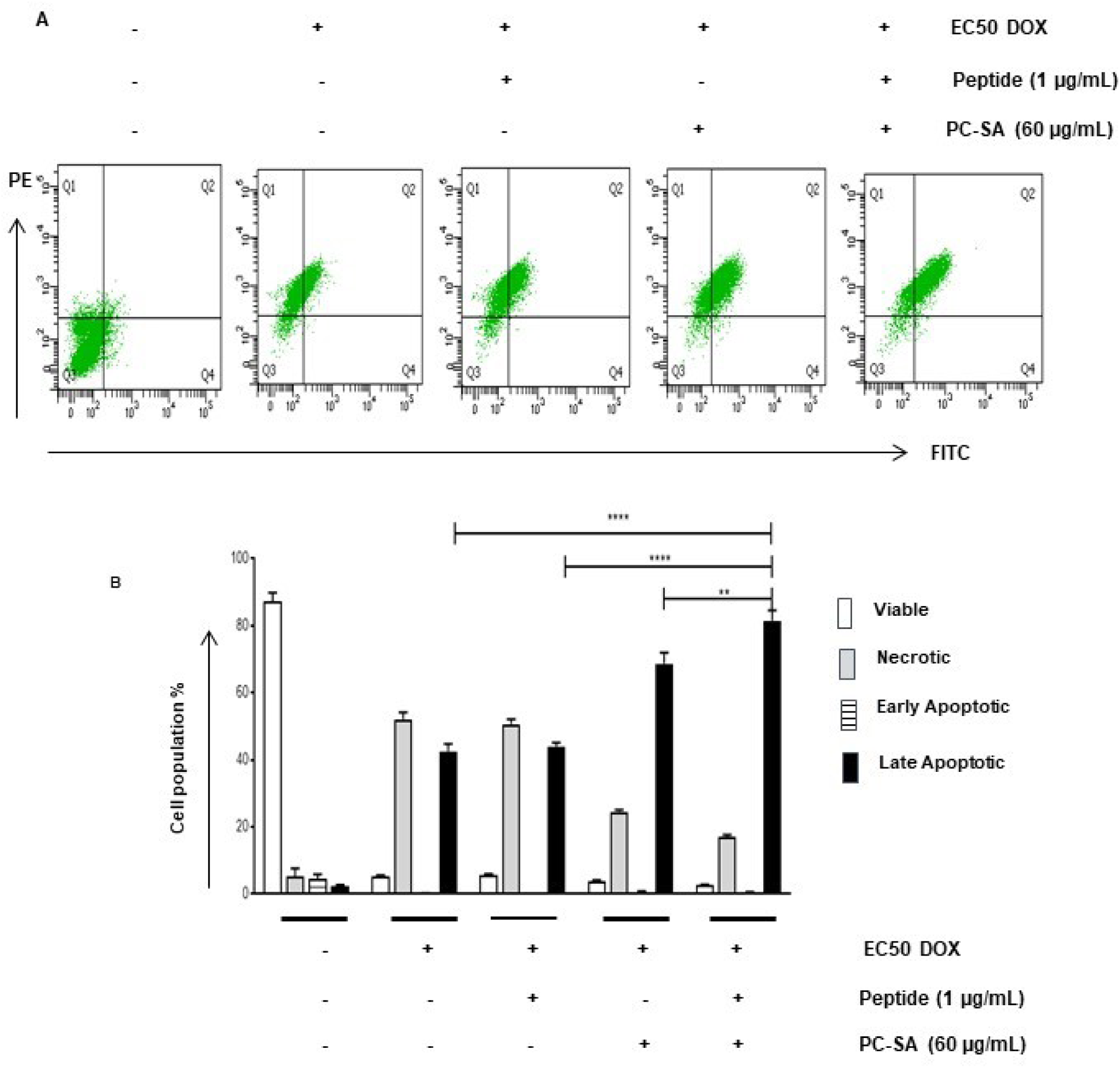
PC-SA:N-PepAc induces apoptosis in SW480 cells. **(A)** Representative dot plots of different treatment groups are shown. **(B)** The graph shows percentage of viable, necrotic, early apoptotic and late apoptotic cell population of different treatment groups at 48 h post EC_50_ DOX addition. Here the used peptide was N-PepAc. Data are representation of three independent experiments. The **, **** represent *p* ≤0.01 and *p* ≤0.0001 respectively.

### PC-SA:N-PepAc downregulated genes associated with tumor progression

Resistance to chemotherapy remains a major problem in cancer-associated death. MDR1 was reported as the most frequently expressed drug resistance gene in cancer cells bearing GOF mutp53 alleles promoting tumor progression (5), (32). PCNA, a marker for abnormal cell proliferation was also found associated with cancer cells bearing p53R273H causing genomic instability (3), (33), (34). SW480 cells were incubated with PC-SA liposome alone or PC-SA:N-PepAc for 2 h followed by incubation with fresh media and treatment with a fixed dose of DOX (0.5µg/ml) for 24 h. Cells incubated with only media and only DOX, N-PepAc along with DOX, PC-SA liposome in addition to DOX for 24 h served as controls. RNA was isolated from different groups of cells, reverse transcribed and cDNAs were subjected to qRT-PCR analysis with GAPDH as the reference internal control gene. We found elevated expression of both MDR1 (Fig. 4A) and PCNA (Fig. 4B) following treatment with DOX in p53R273H bearing SW480 cells. Expression of both the genes was markedly downregulated when cells were pretreated with PC-SA:N-PepAc for 24 h followed by DOX treatment demonstrating the potential of the peptide to reverse enhanced cell proliferation and MDR1-mediated drug resistance in p53R273H expressing cancer cells.

**Fig. 4.**
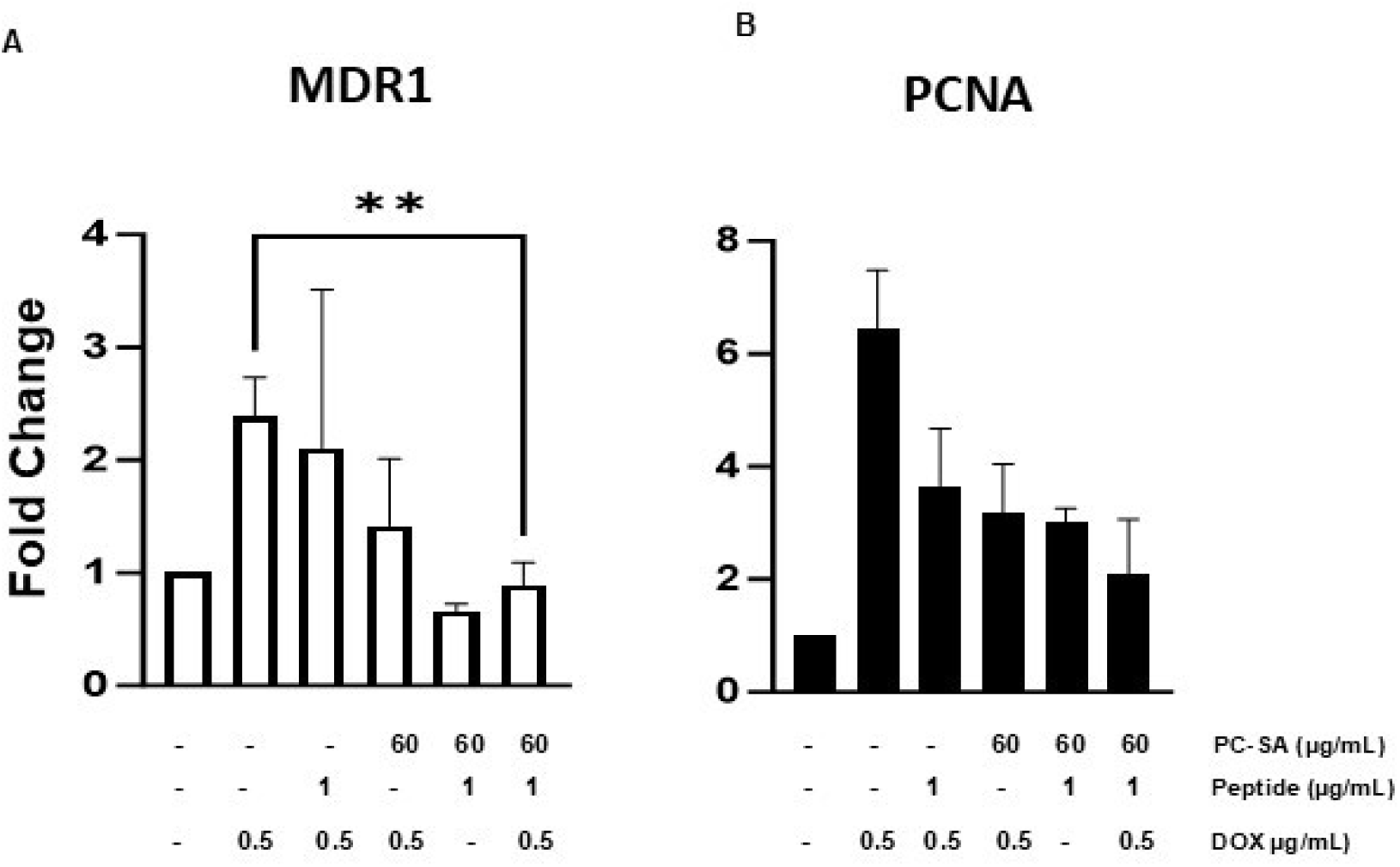
PC-SA:N-PepAc downregulates mutant p53 gain-of-function genes in cells bearing R273H mutation. **(A)** qRT-PCR analysis of MDR1 in SW480 cells treated with indicated groups. (B) qRT-PCR of PCNA of the identical cells treated similarly as A. Data represent fold changes of genes. N-PepAc was the used free and entrapped peptide. All histograms were expressed as Mean ± SD of three independent experiments. ** denotes *p* < 0.01.

### PC-SA:N-PepAc activated apoptosis-inducing genes, caspases and enhanced ROS-activity in p53R273H-bearing cells

We observed significant induction of late apoptotic cell death by PC-SA:N-PepAc. To understand the mechanism of apoptosis we first checked genes that often mediate apoptosis. qRT-PCR analysis revealed marked upregulation of NOXA, PUMA, and particularly BAX by PC-SA:N-PepAc treated group compared to other treatment groups (Fig. 5A). Immunoblot studies confirmed expression of the genes at protein level also (Fig. 5B). As hallmarks of the degradation phase of apoptosis, which include DNA fragmentation, ROS generation, membrane blebbing, and cleavage of several proteins by effector caspases, we checked caspase induction by PC-SA:N-PepAc. Cells sensitized with PC-SA:N-PepAc followed by DOX treatment, demonstrated higher expression of cleaved caspase 3, cleaved caspase 9 and cleaved PARP compared to other groups suggesting a caspase-dependent cell death (Fig. 5C). Preincubation with caspase inhibitor resulted in decreased cell death (Fig. 6A) in all doses of PC-SA:N-PepAc. This proved that DOX-induced cell death observed upon PC-SA:N-PepAc pre-treatment was due to caspase-dependent apoptosis. We further studied PC-SA:N-PepAc pre-treatment followed by DOX induced intracellular ROS generation in drug-resistant cells. By flow cytometry, we observed 80% cell population generated ROS compared to untreated cells (Fig. 6B). Thus, pre-treatment by PC-SA:N-PepAc, a wild-type p53-independent apoptotic pathway is activated by DOX as was previously reported in another study (35).

**Fig. 5.**
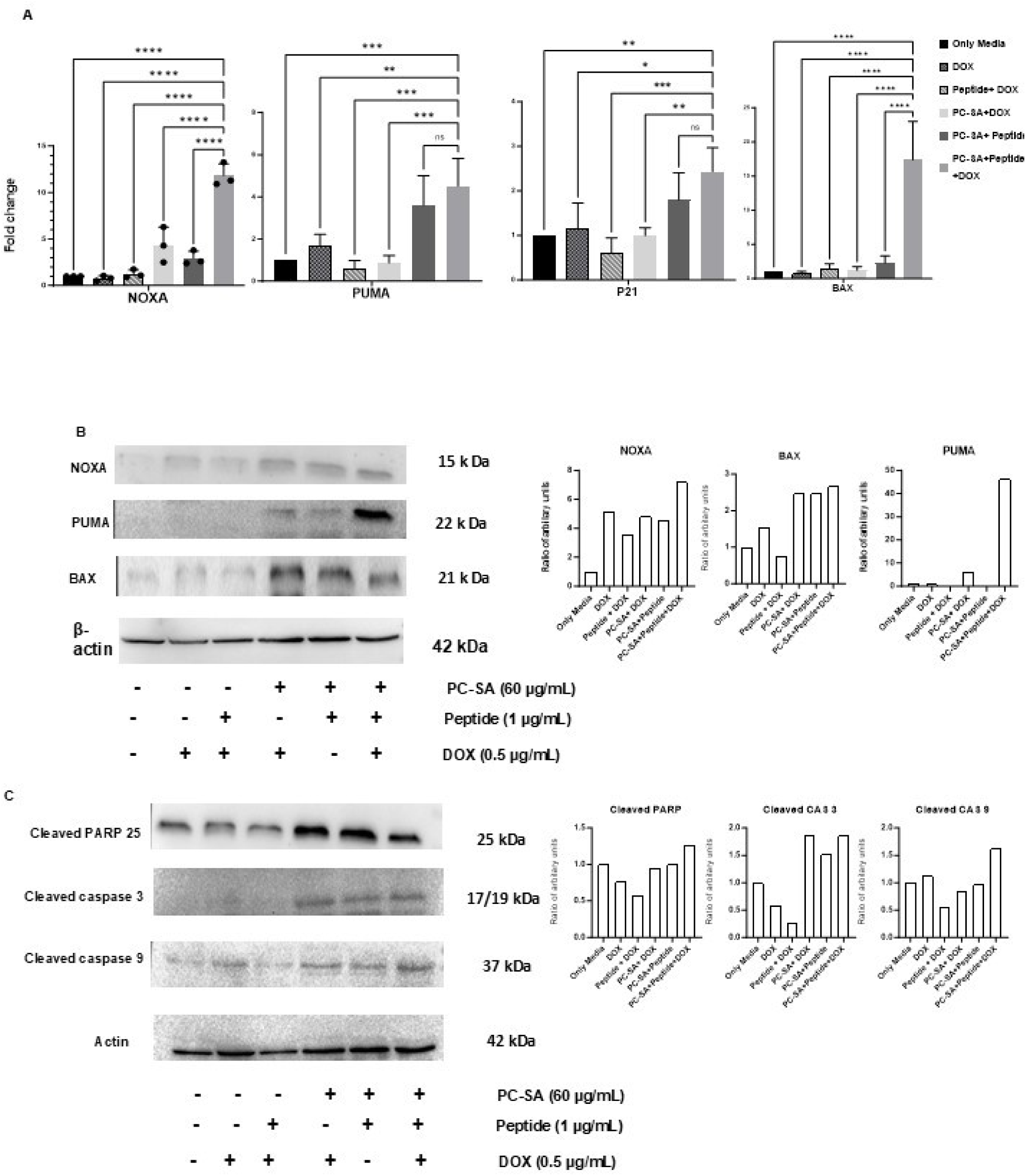
The mechanism for sensitization of cancer cells to DOX by the PC-SA:N-PepAc. **(A)** Expression of NOXA, PUMA, p21 and BAX in indicated treatment groups. **(B)** Western blot showing respective protein levels of NOXA, PUMA and BAX, β-actin expression showed equal protein loading in each well. **(C)** Western blot of cleaved caspase 3, cleaved caspase 9, cleaved PARP. Densitometry of each blot with respect to their loading control is shown next to the blots. All peptides were N-PepAc. Significance is denoted by \**p* < 0.05, ** *p* < 0.01, *** *p* < 0.001, **** *p* < 0.0001.

**Fig. 6.**
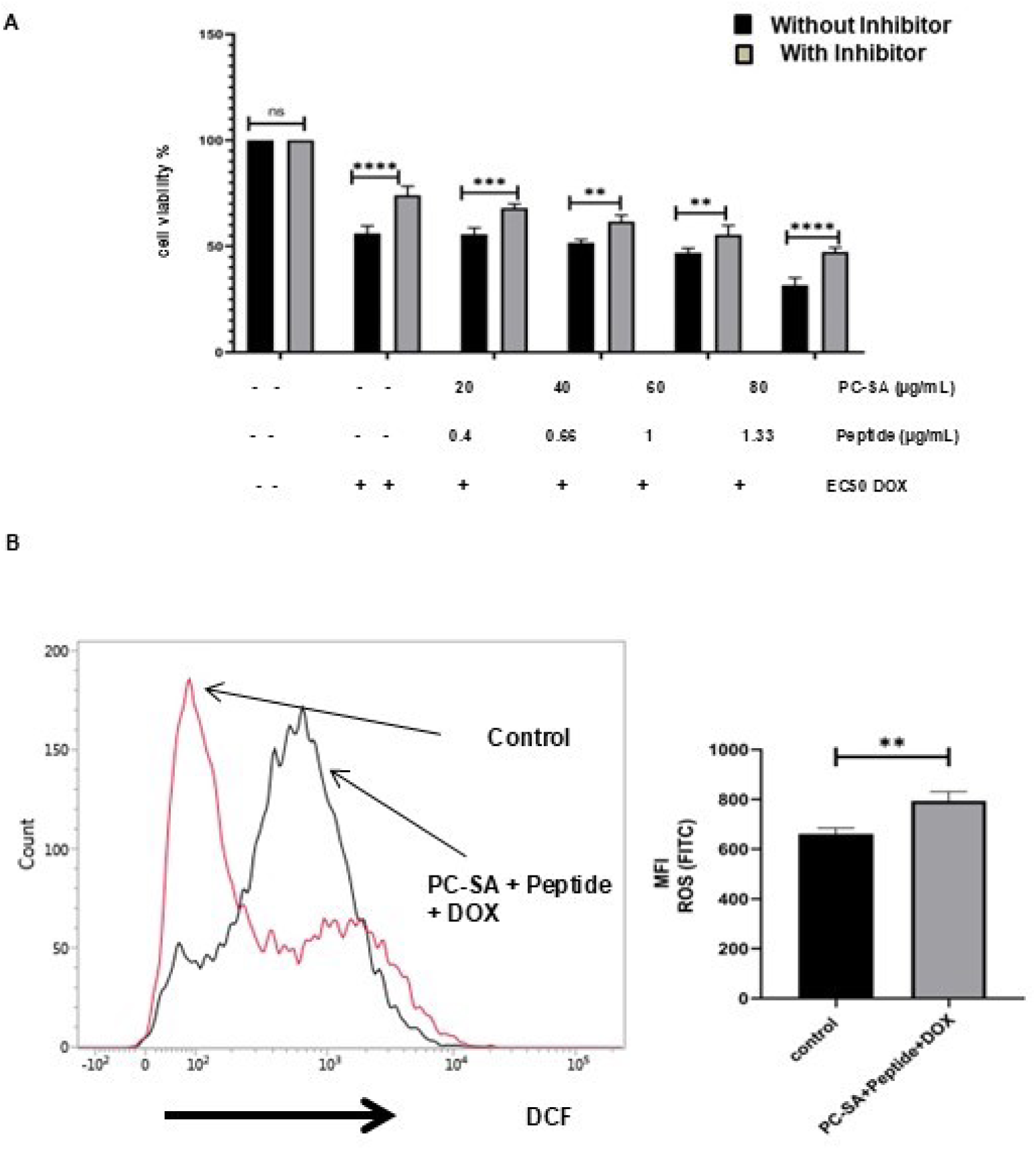
PC-SA:N-PepAc induces cell death through a caspase mediated pathway and ROS generation **(A)** Cell viability was determined following preincubation with caspase inhibitor ZDVQFMK in cells before treatment with DOX and graded doses of liposomal peptide **(B)** Comparison of ROS activities between control and combination treatment (PC-SA:N-PepAc) at 24 h post DOX addition by flow cytometry. Free and entrapped peptides were N-PepAc. Significance is denoted by ** *p* < 0.01, *** *p* < 0.001, **** *p* < 0.0001.

**Fig. 7.**
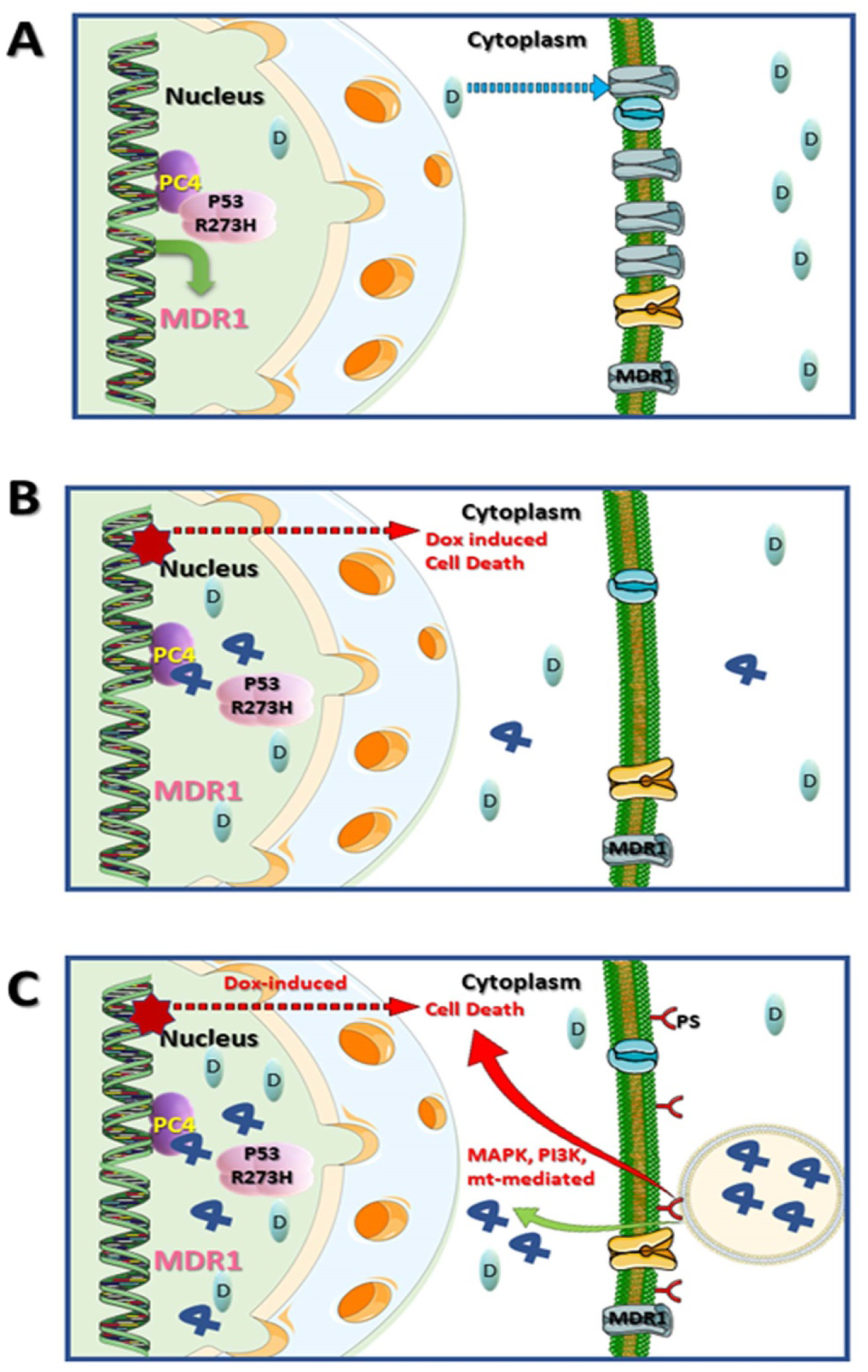
A strategy to target GOF properties of p53R273H. (A) Effect of doxorubicin alone on cells bearing p53R273H. p53R273H binds to its transcriptional coactivator, PC4, and activates the expression of genes responsible for the GOF activity. Activated transcription of the MDR1 gene results in increased MDR1 levels and, consequently, expulsion of doxorubicin, thus leading to chemoresistance. (B) The same cells were pre-treated with R-N-PepAc followed by doxorubicin treatment. Pre-treatment with the peptide disrupts the PC4-p53R273H interaction, leading to a significant reduction in MDR1 transcription. The reduced transcription of the MDR1 gene leads to a substantial reduction in the MDR1 transporter in the membrane, thus causing reduced reflux and retention of doxorubicin in the cell. This leads to DNA damage and cell death. (C) The tumor cells bearing the p53R273H mutation were treated with liposomal NLS-p53(380–386) peptide followed by DOX. The treatment results in the release of N-PepAc into the cancer cells in significant concentrations, which enters the nucleus and blocks the binding of the p53R273H with PC4 by competing with p53R273H for PC4 and thereby abrogates the GOF activities of mutant p53, and doxorubicin-mediated cell death. Additionally, the liposome also induces apoptosis through MAPK, PI3K, and mitochondria-mediated cell death. In the figures, D stands for doxorubicin, PS stands for phosphatidylserine, and the blue ribbon-like object is the peptide.

## Discussion

Specific targeting of GOF-mutp53 still remains a major challenge. Many attempts have been made to target this class of mutant proteins, without any significant success so far (6), (7). It is generally believed that the new gain-of-function in some hotspot mutants of p53 has causal relationship with interaction with some novel protein partners, that is, these protein-protein interactions are necessary for promotion of aggressive properties (36). Once these partner proteins are identified and a causal relationship with gain of new functions is established, these protein-protein interactions become a desirable drug target for abrogation of aggressive gain of function properties. In a previous study, we have shown that interaction of PC4 with a hotspot mutant p53, p53R273H, is necessary for conferring drug resistance to a tumor cell bearing the mutant allele. In that study, a peptide, R-N-PepAc, showed the efficacy of reversing the drug resistance by disrupting this interaction in p53R273H mutant cell lines, thus establishing PC4-R273Hp53 interaction as a drug target in tumors bearing this particular mutation. Thus, it showed promise to target this hotspot mutation (11).

Peptides have shown promise as anticancer therapeutics because of their target specificity, low toxicity and particular capability of selectively targeting protein-protein interaction (12), (13), (37), (38). Although strong anticancer activities have been reported in many cases, their real-life application is limited due to their ease of proteolytic degradation, clearance from plasma, and instability in the gastro-intestinal tract (13). Due to these adverse pharmacokinetic and pharmacodynamic properties, biologically active peptides need a proper delivery system to achieve effective doses in the target tissue. Although the modified peptide, R-N-PepAc, shows good efficacy to reverse many of the gain-of-function properties of a hotspot mutant p53R273H in vitro, its further development as a therapeutic agent requires a proper tissue targeted delivery system. In this study we observed PC-SA:N-PepAc could enhance chemosensitivity to DOX leading to apoptotic cell death of adenocarcinoma cells with p53R273H hotspot mutation. The PC-SA liposome, which has inherent anti-tumor properties, could impart additional cell killing activity that enhances the anti-tumor activity of the peptide-liposome formulation.

N-PepAc at very low dose (1 µg/mL) encapsulated in 60 µg/mL liposome could significantly restore drug sensitivity of two adenocarcinoma cell lines with p53R273H hotspot mutation. The DOX-induced cell death was significantly enhanced when compared to effects shown by the free N-PepAc (*P* < 0.0001), free PC-SA liposome (*P* < 0.001) followed by DOX and free DOX (*P* < 0.0001). Although two free peptides, with and without cell-penetration tag six arginine (R6) could enhance DOX sensitivity to p53R273H bearing cells, they were effective at significantly higher doses (50 and 100 µg/mL). We could reduce the effective dose of free peptide approximately 50-fold when the peptide was entrapped in liposome. PC-SA liposome here served two functions: targeted delivery of peptide to cancer cells as SA has affinity to bind with exposed PS of cancer cells, and anticancer effect of PC-SA liposome itself (17). The restoration of drug sensitivity was further confirmed by reduced expression of drug-resistant genes MDR1 and PCNA by PC-SA:N-PepAc. This formulation could result in 80% cell population to undergo late apoptotic cell death when stimulated with DOX and the drug-induced apoptosis was significantly higher than PC-SA or N-PepAc alone with DOX treatment. We propose that PC-SA:N-PepAc when applied into cell, migrated to nucleus (confirmed in confocal microscopy) assisted by the NLS tag and bound with PC4. This interaction resulted in disruption of PC4-p53R273H interaction and apoptotic cell death upon DOX treatment. Several fold-induction of apoptosis inducing genes (NOXA, PUMA, and BAX), expression of caspases and generation of ROS are linked to programmed cell death pathway, which strongly confirm our proposed mode of action. We also note that this happens in absence of a functional p53, suggesting that DOX can induce p53-independent cell death pathways (35).

Entrapping peptide in a cationic liposome having selective binding to phosphatidyl serine, is a new concept. The formulation controls drug resistance induced by GOF activity of a mutp53 by disrupting an interaction with a partner protein in a mutant p53 expressing cancer cells. The formulation should overcome the bioavailability problems as the peptide is protected from enzymatic degradation by the PC-SA liposome and expected to impart favorable pharmacokinetic properties. The delivery system makes the peptide effective at very low doses, which will reduce the cost and toxicity of the free peptide. The property of PC-SA liposomes to specifically target cancer cells, sparing the healthy cells, addresses the challenge of targeting a protein-protein interaction specifically, a major concern of cancer therapy. In addition, antitumor effects of the PC-SA liposome itself induces combined antitumor effects in association with the N-PepAc peptide. We speculate that such dual action may reduce the chance of acquiring other mechanisms of drug resistance. It may be possible to target other protein-protein interactions with PC-SA liposome entrapped peptides.

## Conclusion

We described a peptide-liposome formulation that can act on a particular GOF-mutantTP53 bearing cell to abrogate the GOF activity and enhance cell killing. This formulation may offer a solution for both a delivery and dual cell killing mechanisms.

## Abbreviations and Glossary

PC4,: positive cofactor 4
DOX,: doxorubicin;
DBD,: the DNA-binding domain (residues 102 to 293) which plays important role in DNA binding by p53;
R273H,: arginine at 273 position is mutated to histidine
R248Q,: arginine is mutated to glutamine at 248 residue
R175H,: arginine at 175 position is mutated to histidine
R249S,: arginine mutated to serine
GOF,: gain of function
LOF,: loss of function; mutp53, mutant p53
PC-SA,: phosphatidylcholine-stearylamine
MDR1,: multidrug-resistant gene; PCNA, proliferating cell nuclear antigen
PepAc,: p53(380-386, 3Ac), a triply lysine acetylated (K381, K382, and K386) peptide encompassing the residues 380-386 of the p53 protein
N-PepAc,: NLS-p53(380-386, 3Ac), PepAc with an attached nuclear localization signal
RN-PepAC,: R6-NLS-p53(380-386, 3Ac), PepAc attached to a cell penetrating sequence (six D-arginine residues, R6) and nuclear localization signal sequence (NLS)
PC-SA:N-PepAc: N-PepAc entrapped in PC-SA liposomes.

## Acknowledgement

The infrastructure support of Bose Institute, Kolkata for peptide synthesis and CSIR-Indian Institute of Chemical Biology for cell culture work is highly acknowledged.

