## supplementary figure for "A Dual-Action Liposome-Peptide Formulation Synergistically Counteracts A Gain-of-Function p53 Mutant"

### Gating strategy

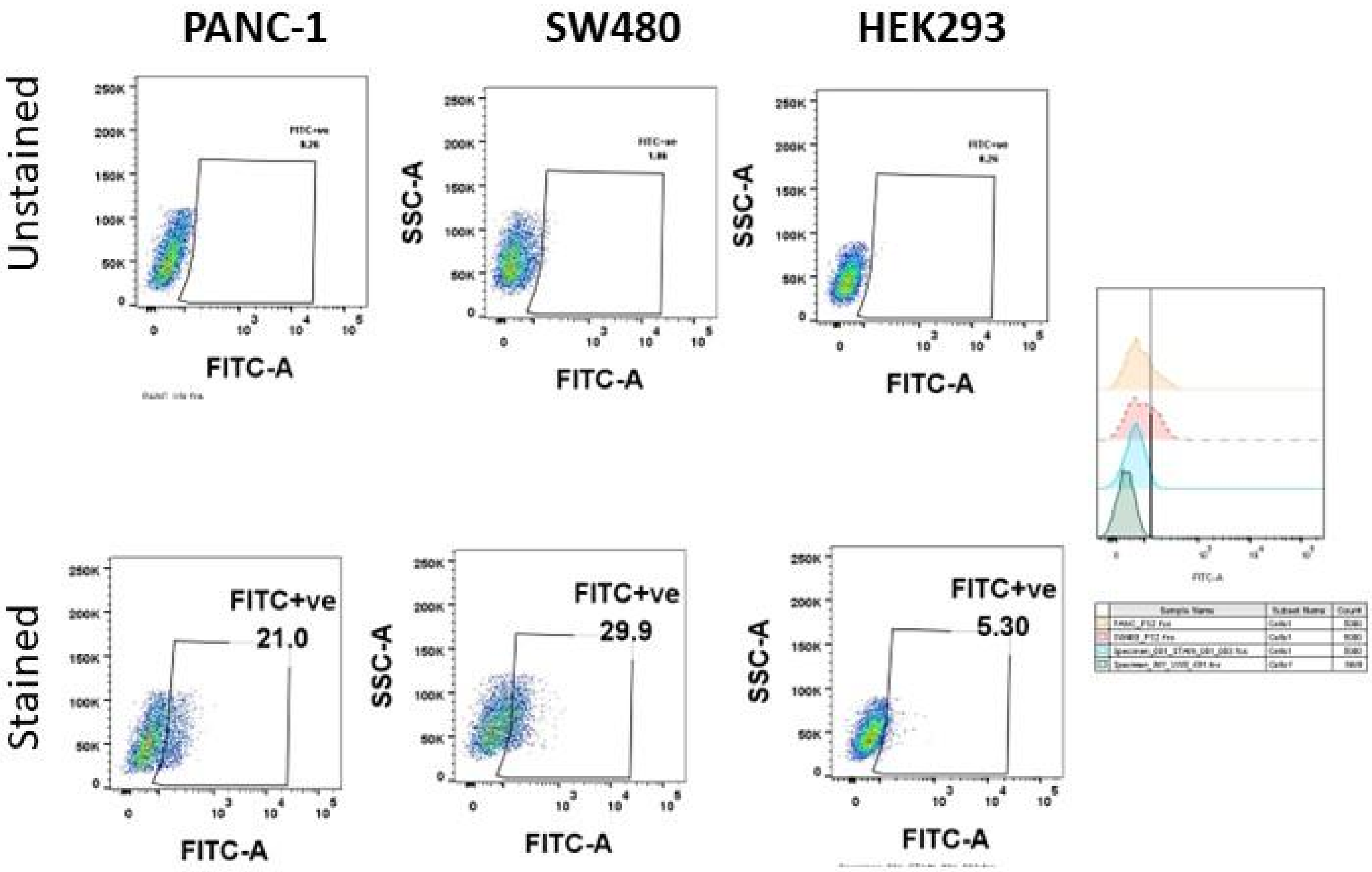


Figure S1 Gating strategy for estimation of surface exposed phosphatidylserine: PANC-1, SW480 and HEK 293 cells were harvested and stained for FITC Annexin-V antibody, ten thousand events recorded on flow. The dot plots here represent the gating strategy for positive PS population which were set based on the unstained cells in each experiment. The gates were set same for all samples.
